# Ecological succession revisited from a temporal beta-diversity perspective

**DOI:** 10.1101/2022.09.11.507494

**Authors:** Ryosuke Nakadai, Satoshi N. Suzuki

## Abstract

1. Ecological succession, which is the community re-assembly process after a disturbance, is a study topic receiving renewed attention in relation to anthropogenic disturbance as well as one of the most classical ones in ecology. Previous studies occasionally revealed that compositional shifts decrease toward late succession stages and discussed the potential links with species life history and longevity. However, clear empirical evidence is not available until now because of limited analytical approaches. Therefore, traditional approaches used in previous studies could not quantify the relative contribution of demographic processes to apparent compositional shifts in communities.
2. In the present study, we aimed to understand ecological succession processes by revealing the patterns of temporal beta diversity based on both conventional Bray-Curtis dissimilarity and recently developed individual-based dissimilarity indices using a long-term dataset. Specifically, we used published forest inventory data from permanent forest plots in cool temperate forests along a secondary successional chronosequence, with stands at 17 to 106 years post clear-cutting.
3. We clearly demonstrated the detailed patterns of temporal beta-diversity indices (i.e., conventional Bray–Curtis dissimilarity and individual-based dissimilarity indices) based on stem number and stem basal area along long-term chronosequences across approximately one hundred years.
4. *Synthesis*. Using a long-term forest inventory dataset, this study demonstrated the link between the apparent compositional shifts and the changes in each component of the demographic processes (i.e., recruitment, growth, and mortality) during secondary forest succession in the context of temporal beta diversity. As done in this study, future research on changes in community composition during ecological successions at various sites and systems will help elucidate the relationships between temporal changes in global biodiversity and the impact of anthropogenic environmental changes.

## Introduction

Ecological succession, the processes of re-assembly and change over time following natural or anthropogenic disturbance, is the most classic theme in ecology (Clements 1916), yet one that has received renewed attention recently (Prach and Walker 2011, Chang and Turner 2019). Ecological succession is generally driven by the competition-colonisation trade-off (Rees et al., 2001) in dominance-controlled communities (Yodzis, 1986) and has been reported in various taxonomic groups (Hill et al., 2002; Pascual-García & Bell, 2020). Regardless of taxonomic group, previous studies revealed that diversity is low in the early stages when several pioneer species are introduced, and reaches a maximum in the mid-succession period when mid-succession and climax species co-occur, and finally declines due to competitive exclusion among climax species (Begon & Townsend, 2020). Most of the studies employed the space-for-time substitution approach (i.e., the chronosequence approach) as an alternative to long-term studies (Pickett, 1989; Suzuki & Hirao, 2018). Consequently, revealing patterns of actual compositional shifts over time is still an important missing piece for understanding the processes of ecological succession and more generally, community assembly.

Temporal changes in community composition are referred to as temporal beta diversity (Legendre, 2019) and have recently been well studied (Gotelli et al., 2022; Brice et al., 2019) given the growing concerns regarding the global biodiversity crisis (Magurran et al., 2019). Recent studies have attempted to elucidate the background processes of temporal compositional shifts by developing various kinds of statistical approaches (Gotelli et al., 2022; Legendre, 2019; Shimadzu et al., 2015; Tatsumi et al., 2021). Among these trends in the temporal beta-diversity field, the directional successional changes have been found to greatly affect the observed temporal beta diversity (Li et al., 2016; Magurran et al., 2019). Therefore, revealing the relationships between temporal beta diversity and ecological succession would have a great impact on future studies focusing on biodiversity changes.

Temporal beta diversity along an ecological succession can be referred to as succession rate (Foster & Tilman, 2000; Chazdon et al., 2007; Li et al., 2016). Succession rate is thought to be related to differences in growth rate, life history, and longevity among species (Foster & Tilman, 2000). These species differences can affect the succession rate through demographic processes of individuals (i.e., recruitment, growth, and mortality). However, clear empirical evidence is currently lacking because of analytical limitations. Consequently, traditional beta-diversity indices could not quantify the relative contribution of the demographic processes to apparent community compositional shifts found in snapshot datasets. In order to better quantify temporal beta diversity, Nakadai (2020) developed an extended approach to harmoniously include information on both the individual persistence and the turnover in compositional shifts in a community; this approach was named “individual-based temporal beta diversity” (Nakadai, 2020, 2021, 2022). Specifically, the approach can connect the individual demographic processes with apparent compositional shifts in a community over time (Nakadai, 2020, 2022). This can strengthen the understanding of the background processes causing temporal compositional shifts, which cannot be revealed using only the relevant species information and species abundance data. Therefore, it is important to describe patterns of ecological succession that can affect patterns of temporal beta diversity based on relevant species information and species abundance data, and to assess the impact of the processes behind these patterns by using individual-based beta-diversity indices. In the present study, we revisited the classical issues of ecological succession from the perspective of temporal beta diversity and aimed to understand the processes of ecological succession by revealing the patterns of multiple types of temporal beta-diversity indices.

For this purpose, we used long-term forest inventory data in cool temperate forests along a secondary successional chronosequence of stands between 17 and 106 years after clear-cutting. In a typical forest stand development, after a stand-replacing disturbance (i.e., secondary succession), developmental stages can have four distinct phases, namely stand initiation, stem exclusion, understory re-initiation, and old-growth (Oliver & Larson, 1996). At the initiation phase, a number of pioneer trees recruit first, followed by mid- and late-successional species. Therefore, species composition based on stem density is expected to change drastically during the early stage, which would be driven by the high rate of recruitment. When species composition is evaluated based on dominance (i.e., stem basal area), differences in stem growth between species also contributes to the compositional shift because growth rate greatly varies between species in this phase. As the stand develops, competition between trees becomes intense, resulting in exclusion of the stems of suppressed trees. In the stem exclusion phase, recruitment rarely occurs because of the complete closure of the canopy. Therefore, compositional shifts based on stem density will be driven by mortality during the mid-stage. Stem growth will also contribute to the compositional shift based on dominance at this stage. After the stem exclusion phase, as the stand matures, individual turnover rate becomes slower. However, large tree mortalities occur occasionally, creating canopy gaps, with stem recruitment occurring in these gaps. In the understory re-initiation phase and subsequent old-growth phase, because a stronger filtering process will occur through gap regeneration than through mortality (Suzuki & Hirao, 2018), compositional shifts based on stem density will be driven by recruitment. In contrast, compositional shift based on dominance would be driven by the mortality of large trees and not by recruitment. Specifically, we focused on the following four hypotheses: (1) the degree of compositional shift decreases toward the late stage of ecological succession; (2) the relative speed of compositional shift relative to individual turnover is consistent during the progression of ecological succession, thus both turnovers of stem number and stem basal area primarily drive apparent compositional shift; (3) based on stem number, compositional shift is primarily driven by the recruitment of individuals in the early and late stages of the succession and mortality in the middle stage; and (4) based on stem basal area, stem growth mainly drives the compositional shift, but the relative contribution of mortality increases as ecological succession progresses.

## Materials and methods

### Study site and census data

We used published forest inventory data from permanent forest plots in secondary forests in the University of Tokyo Chichibu Forest (35°53′–58′ N, 138°46′–139°0′ E) located in the Oku-Chichibu mountains of central Japan (Igarashi et al., 2005; Saito et al., 2014; Saiki et al., 2019). In 1982 or 1987, 15 permanent plots were established in cool temperate secondary forests at 780–1200 m elevation (Table S1, Fig S1). The plot areas ranged from 0.08 ha to 0.15 ha. Stand ages, calculated as years after clear-cutting, ranged from 17 to 81 years at the time of plot establishment. All stems with a diameter at breast height (DBH) larger than 4 cm were tagged and their species were recorded. The DBH was measured for almost all stems in the census, however, some individuals were not measured owing to measurement difficulty (e.g., standing in dangerous positions: 30/16508 time-individual records; 0.0017%). Measurements were carried out every 5 years until 1992 for all plots. After 1992, measurements were collected every 5 years for eight and every 10 years for seven of the 15 plots (1982, 1987, 1992, 1997, 2002, 2007, 2012, and 2017; Fig. 1). At the time of the final census in 2017, the oldest stand was 106 years old; therefore, this data series covers tree stands aged between 17 and 106 years old (Table S1). These plots are the subject of one of the longest running monitoring programs on the impact of anthropogenic disturbance in the world, and the patterns of shade tolerance and above ground biomass have been reported in previous studies (Suzuki & Hirao, 2018; Suzuki, 2021). Therefore, these plots satisfy the requirements for both the chronosequence approach and long-term monitoring. In addition, Foster and Tilman (2000) conducted a follow-up survey at the sites using the chronosequence approach, following the study by Inouye et al. (1987); moreover, this follow-up survey was named “chronosequence resampling”. In this context, the surveys for the plots in the present study were simply named “long-term and multiple chronosequence resampling.” This was done as the name represents one of the most appropriate methodologies for revealing patterns of temporal beta diversity, more specifically, it is an extended procedure of both the chronosequence approach and chronosequence resampling. We limited our analysis to stems with DBH ≥ 5 cm for statistical analysis, because stems that grew larger than 5 cm were treated as recruited stems. We focused only on minimum survey intervals and finally, we could calculate the temporal beta-diversity indices for 81 survey intervals from the 15 plots (Fig. S1, Table S2).

**Figure 1.**
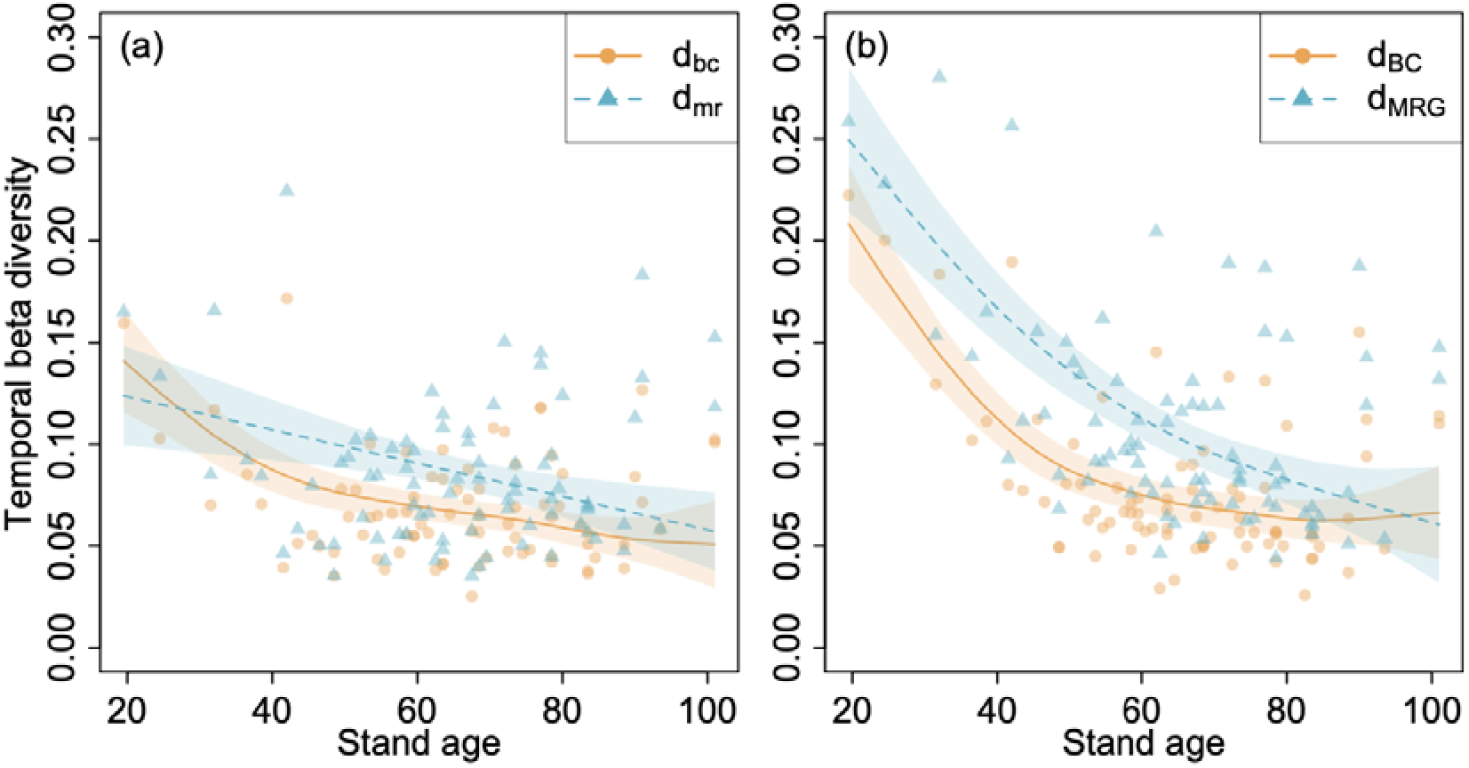
Relationship between temporal beta-diversity indices and stand age of forest plots situated in a cool temperate secondary forest in Japan. Subplots (a) and (b) depict the indices based on stem number (*d*_*bc*_ and *d*_*mr*_) and basal area (*d*_*BC*_ and *d*_*MRG*_), respectively. Each line was drawn in a situation where all values except stand age were fixed to the mean values for each index. Each coloured area represents the 95% confidence intervals for each index. Model structures and selection details are shown in Table 1 and S5.

### Calculating the indices of temporal beta diversity

We focused on two types of Bray–Curtis dissimilarity indices (Odum, 1950) based on both stem number (*d*_*bc*_) and stem basal area (*d*_*BC*_). Hereafter, upper-case and lower-case subscript letters correspond to indices based on stem number and stem basal area, respectively. Bray–Curtis dissimilarity indices can be partitioned into the loss (*d*_*b*_, *d*_*B*_) and gain (*d*_*c*_, *d*_*C*_) components (Legendre, 2019). We also used individual-based temporal beta-diversity indices based on both stem number (*d*_*mr*_; Nakadai, 2020) and stem basal area (*d*_*MRG*_; Nakadai, 2022). When considering individual-based temporal beta diversity, the indices proposed by Nakadai (2020) take only the presence or absence of individuals (i.e., qualitative information) into account, while the indices proposed by Nakadai (2022) can also include the basal area of each individual (i.e., quantitative information). These indices are extensions of the Bray-Curtis dissimilarity index to individual turnover. The individual-based indices linked to stem number can be partitioned into the mortality and recruitment components (*d*_*m*_, *d*_*r*_; Nakadai, 2020), and the indices relating to stem basal area can be partitioned into the growth component, in addition to mortality and recruitment components (*d*_*G*_, *d*_*M*_, *d*_*R*_; Nakadai, 2022). The relative speeds of apparent compositional shifts relative to turnover of stem number and stem basal area were calculated for each survey interval (*v*_*s_QL*_, *v*_*s_QT*_; Nakadai, 2020, 2022). By multiplying the relative speed indices (*v*_*s_mr*_, *v*_*s_MRG*_) by the indices of individual turnover (*d*_*mr*_, *d*_*MRG*_), we can obtain indices of compositional shift due to individual turnover (*v*_*s_mr*_, *v*_*s_MRG*_). Similarly, actual contributions of individual demography (i.e., mortality, recruitment, and growth) to apparent compositional shifts were calculated (*v*_*s_m*_, *v*_*s_r*_, *v*_*s_M*_, *v*_*s_R*_, *v*_*s_G*_). The index *v*_*s_mr*_ was identical to the component of compositional shift (*d*_*s*_=(b+c)/(2p+m+r)) introduced in Nakadai (2020). Note that *d*_*bc*_ and *d*_*BC*_ were completely correlated with *v*_*s_mr*_ and *v*_*s_MRG*_. All the temporal beta-diversity indices used in the present study are summarised in Table S3. We removed the individual for which DBH was missing using a pairwise deletion when the indices of temporal beta diversity were calculated. Pairwise deletion is a standard method employed to calculate genetic distances (Fregin et al., 2012).

**Table 1.**
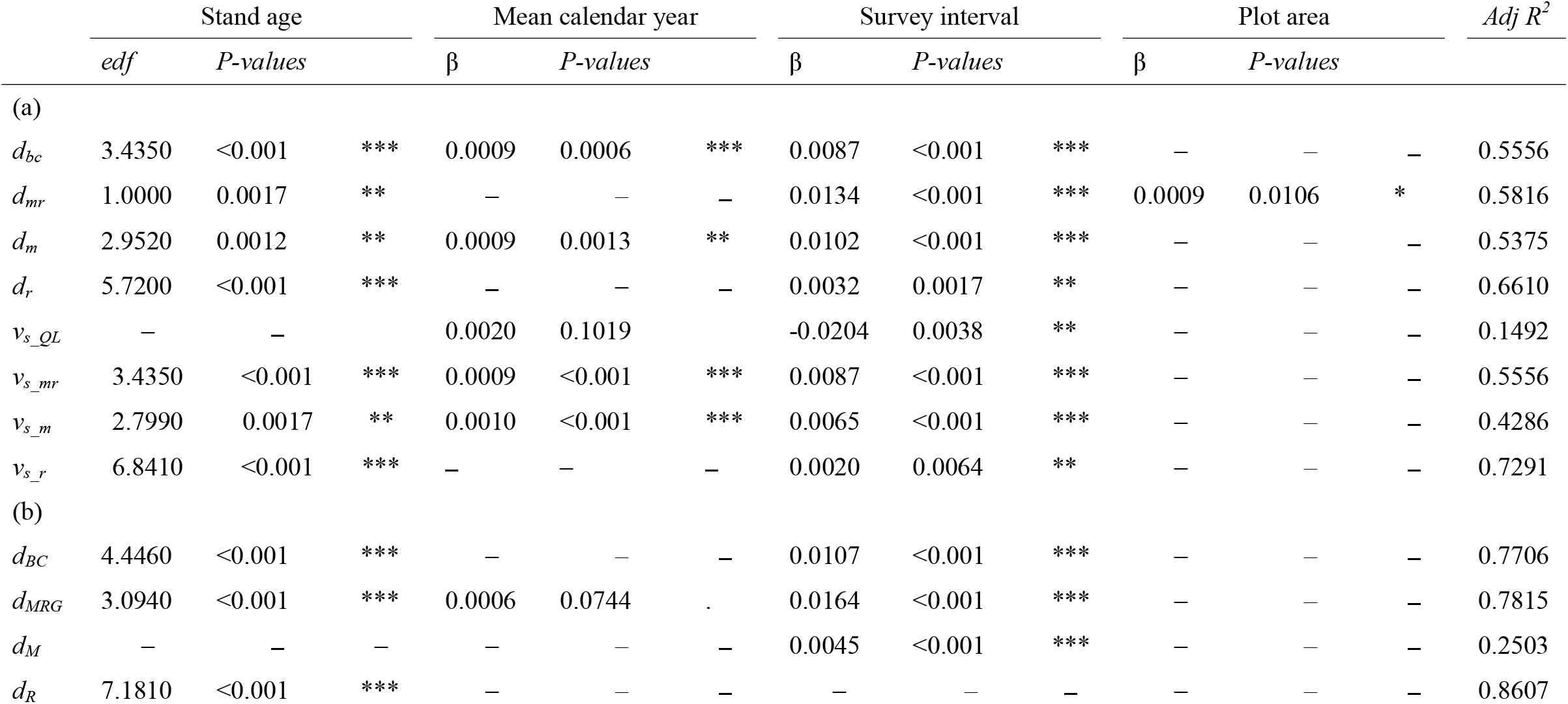
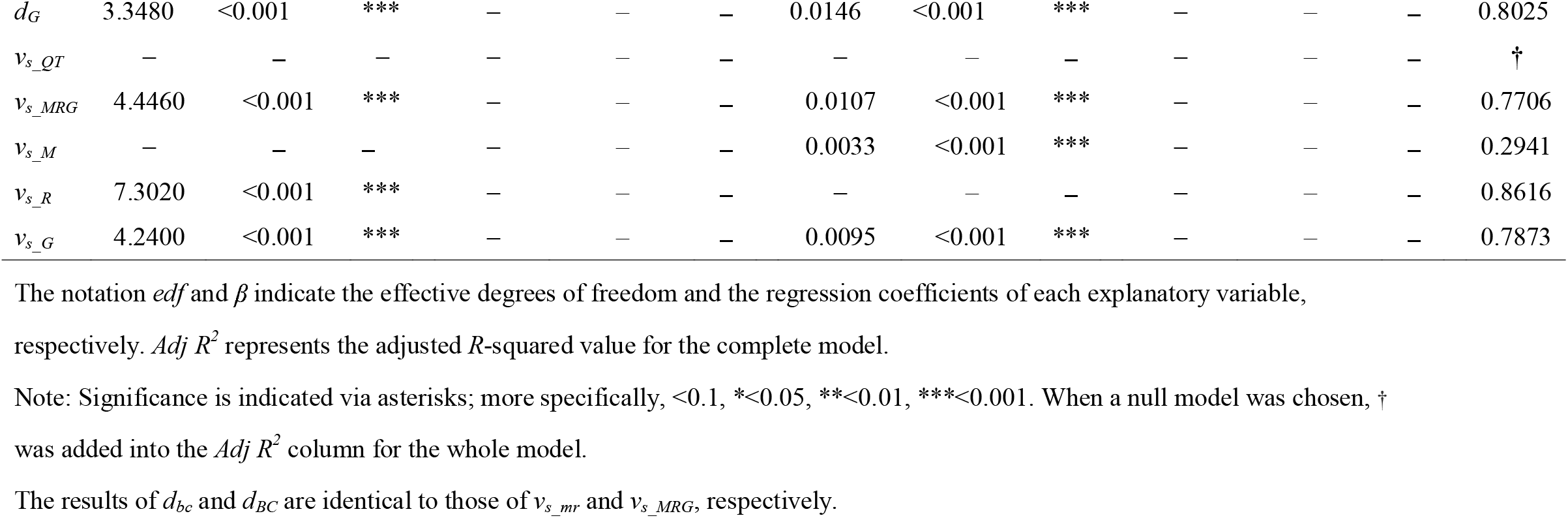
Results of a generalised additive mixed model based on the best models selected using an Akaike information criterion (AIC) comparison for temporal beta-diversity indices based on (a) stem number and (b) stem basal area in established plots in a temperate forest in Japan.

### Statistical analysis

We used generalised additive mixed models (GAMMs) to reveal the relationship of temporal beta-diversity indices with stand age. In addition to stand age, mean calendar year, survey interval, and plot area were considered as explanatory variables, whereas plot identity was considered a random effect. The influence of mean calendar year can be interpreted as anthropogenic environmental changes since the mid-20^th^ century. Both survey interval and plot area were considered additional factors potentially affecting the temporal beta-diversity indices. A smooth spline fit was used for stand age only, whereas linear fits were used for the other explanatory variables in the GAMMs. For each temporal beta-diversity index, model selection was conducted using Akaike information criteria (AIC).

All analyses were conducted using R software (ver. 4.1.0; R Development Core Team, 2021). The R packages *mgcv* (Wood, 2017) and *MuMIn* (Barton, 2022) were used to fit the GAMMs and perform model comparisons using AIC.

## Results

Given the AIC comparison results, the Bray-Curtis dissimilarity indices based on both stem number (*d*_*bc*_) and stem basal area (*d*_*BC*_) significantly decreased with increasing stand age (Fig. 1, Table 1), thus, supporting hypothesis 1. The relative speed of compositional shifts against both stem number turnover (*v*_*s_QL*_) and stem basal area (*v*_*s_QT*_) did not show significant patterns against stand age (Fig. 2, Table 1). These results suggest that contributions of the stem turnover to apparent compositional shifts were constant along the succession. However, the total turnover of both stem number (*d*_*mr*_) and stem basal area (*d*_*MRG*_), as well as their contribution to apparent compositional shifts (*v*_*s_mr*_, *v*_*s_MRG*_) significantly decreased with increasing stand age (Fig. 1ab, Table 1ab), thus supporting hypothesis 2. For the indices based on stem number, the components of recruitment (*d*_*r*_) and their contributions to apparent compositional shifts (*v*_*s_r*_) were large in the early and late stages of the succession, whereas those related to mortality (*d*_*m*_, *v*_*s_m*_) were large in the middle stage (Fig. 3a, 4a), thus supporting hypothesis 3. For the indices based on stem basal area, the components of stem growth (*d*_*G*_) and their contributions to apparent compositional shifts (*v*_*s_G*_) were found to drive the total pattern (*d*_*MRG*_, *v*_*s_MRG*_), however, the relative contribution of mortality (*d*_*M*_, *v*_*s_M*_) increased with an increasing stand age (Fig. 3b, 4b), thus, supporting hypothesis 4. Furthermore, mean calendar year significantly influenced the Bray-Curtis dissimilarity index based on stem number (*d*_*bc*_, Fig. 1, Table 1), as well as the mortality component of total stem number turnover (*d*_*m*_, Fig. 3, Table 1), the contribution of mortality and total stem basal turnover to apparent compositional shifts (*v*_*s_m*_ and *v*_*s_mr*_, Fig. 4, Table 1), and the loss component of the Bray-Curtis dissimilarity index (*d*_*b*_, Fig. S2, Table S4).

**Figure 2.**
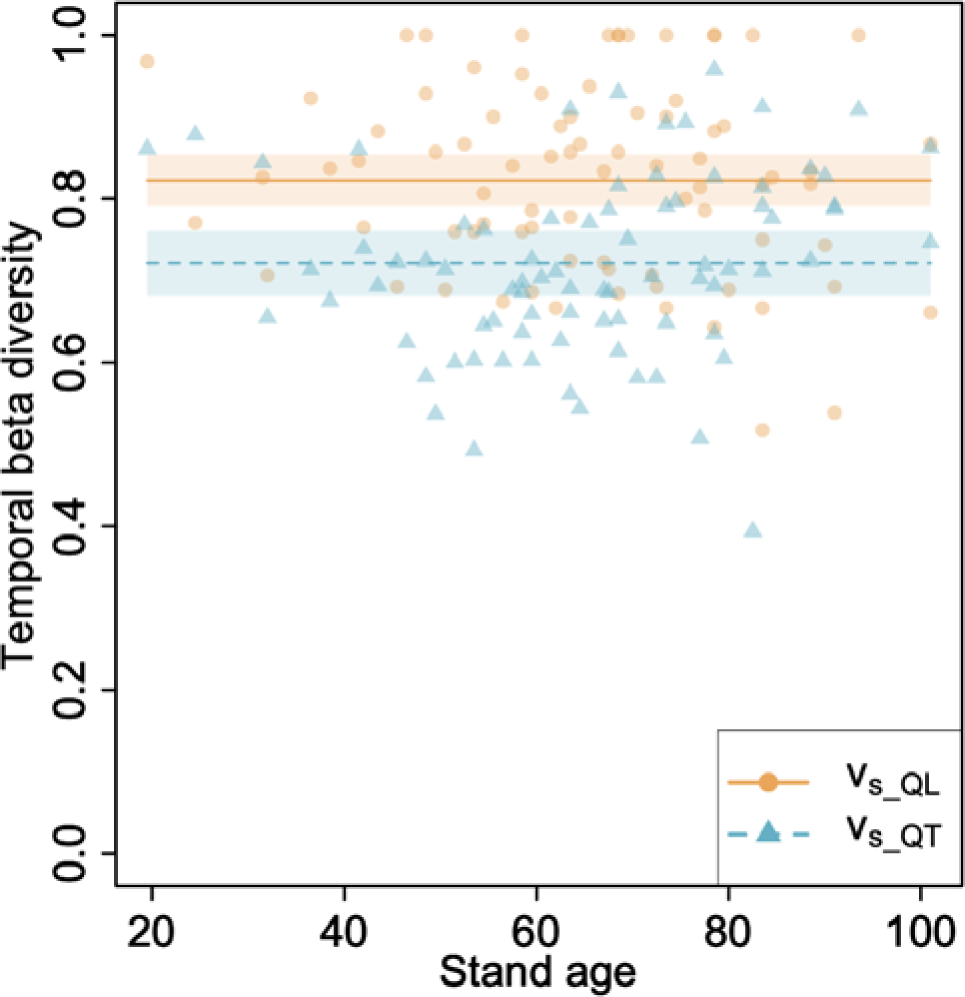
Relationship between the relative speed of compositional shifts relative to individual turnover based on stem number (*v*_*s_QL*_) and basal area (*v*_*s_QT*_), and to the stand age of forest plots situated in a cool temperate secondary forest in Japan. Each line was drawn in a situation where all values except stand age were fixed to the mean values for each index. Each coloured area represents the 95% confidence intervals for each index. Model structure and selection details are shown in Table 1 and S5.

**Figure 3.**
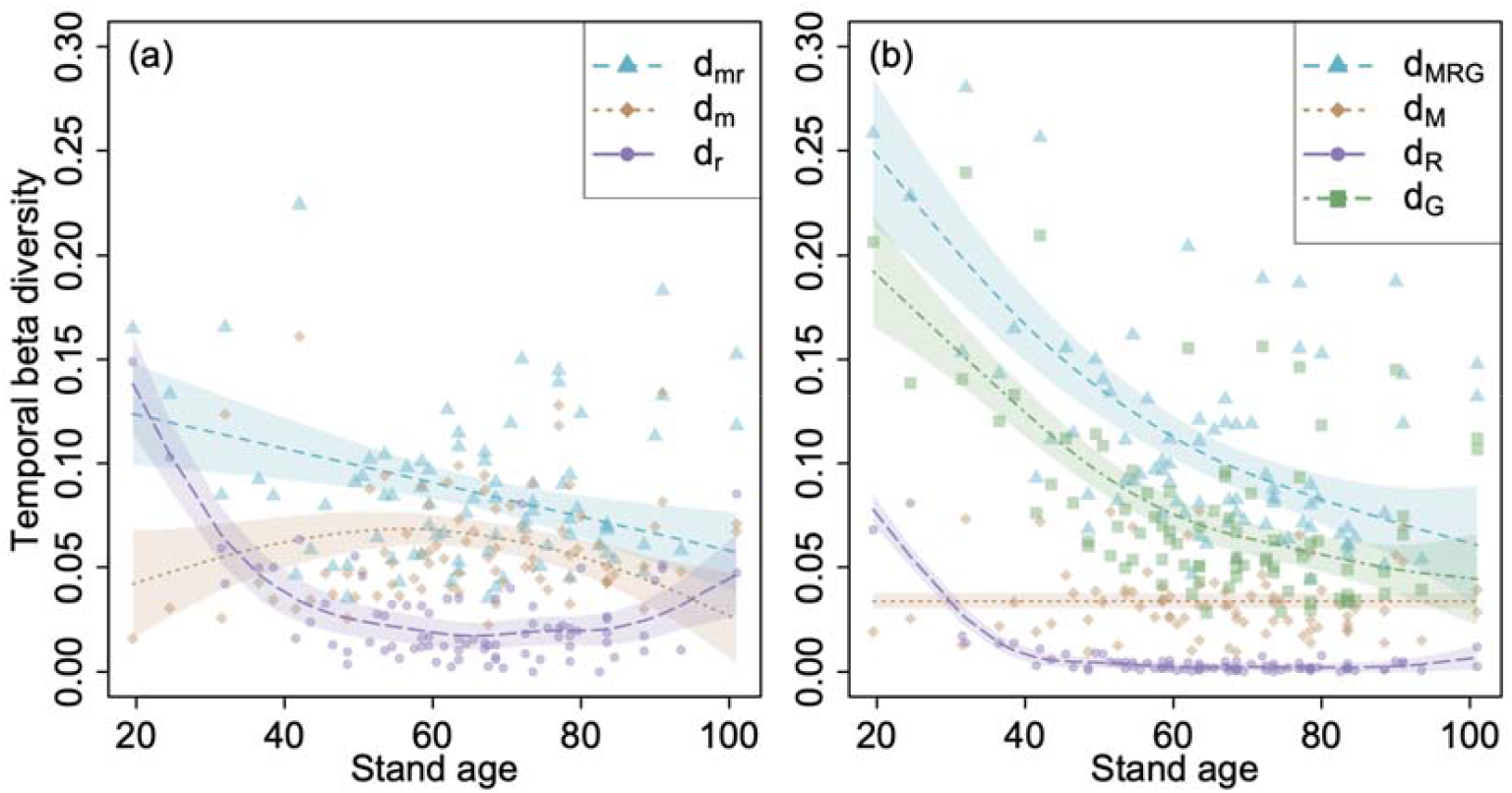
Relationship between each component (i.e., mortality, recruitment, and growth) of individual-based temporal beta-diversity indices and stand age of forest plots situated in a cool temperate secondary forest in Japan. Subplots (a) and (b) depict the indices based on stem number (*d*_*mr*,_ *d*_*m*_, and *d*_*r*_) and basal area (*d*_*MRG*_, *d*_*M*_, *d*_*R*_, and *d*_*G*_), respectively. Each line was drawn in a situation where all values except stand age were fixed to the mean values for each index. Each coloured area represents the 95% confidence intervals for each index. Model structures and selection details are shown in Table 1 and S5.

**Figure 4.**
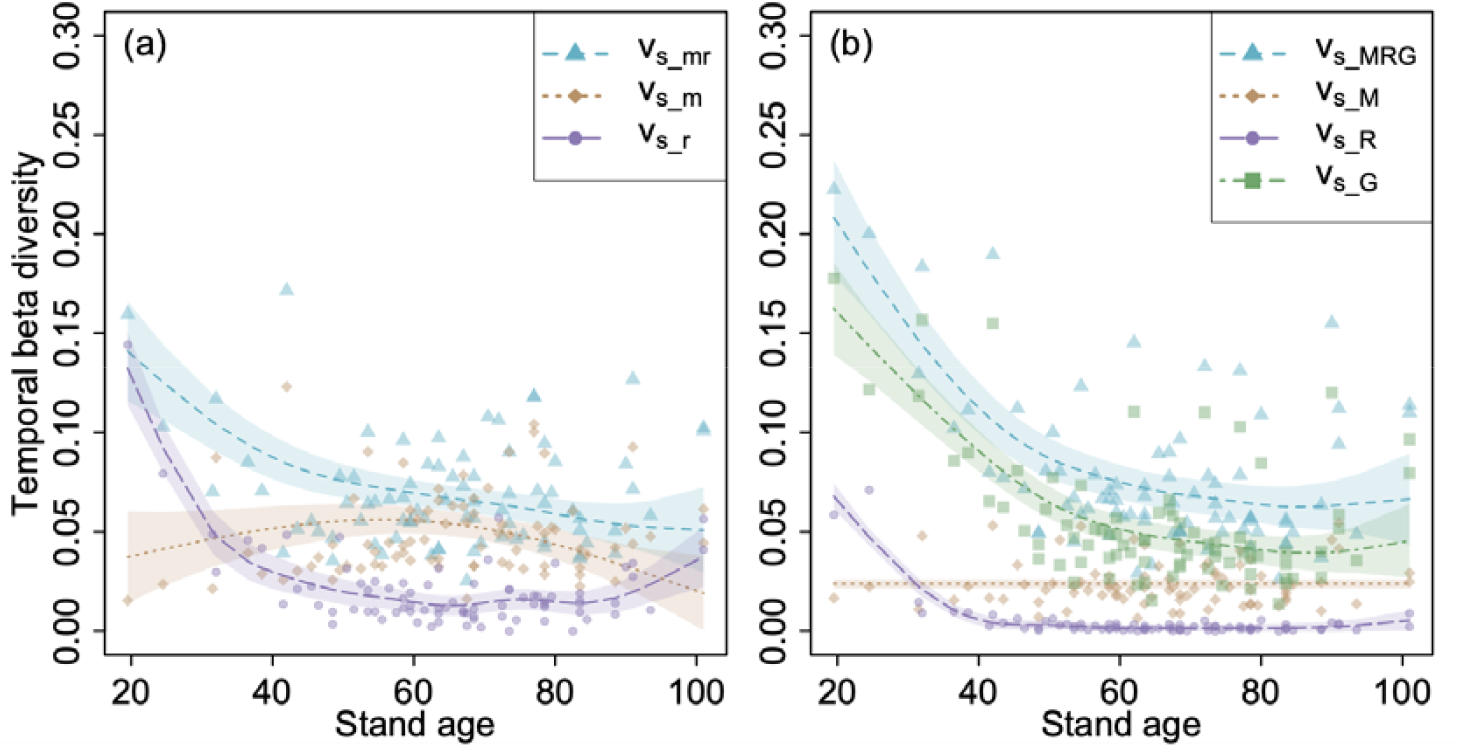
Relationship between each component (i.e., mortality, recruitment, and growth) of individual-based temporal beta-diversity indices and stand age of forest plots situated in a cool temperate secondary forest in Japan. Subplots (a) and (b) depict the indices based on stem number (*v*_*s_mr*,_ *v*_*s_m*_, and *v*_*s_r*_) and basal area (*v*_*s_MRG*_, *v*_*s_M*_, *v*_*s_R*_, and *v*_*s_G*_), respectively. Each line was drawn in a situation where all values except stand age were fixed to the mean values for each index. Each coloured area represents the 95% confidence intervals for each index. Model structures and selection details are shown in Table 1 and S5.

## Discussion

Here we present the long-term, detailed patterns of Bray–Curtis dissimilarity indices based on both stem number and stem basal area, as well as individual-based indices based on stem number and stem basal area, on temporal beta diversity using a long-term and multiple chronosequence resampling across approximately one hundred years. Furthermore, the results presented here support all four of our initial hypotheses. Firstly, we show that the degree of compositional shift decreases toward the later stages of ecological succession (Fig. 1, Table 1). Secondly, the relative speed of compositional shifts relative to individual turnover is shown to be consistent during the progression of ecological succession, thus both stem number and stem basal area turnover primarily drive apparent compositional shifts (Fig. 1, 2, Table 1). Thirdly, based on stem number, compositional shifts were shown to be primarily driven by the recruitment of individuals in the early and late stages of the succession, as well as mortality in the middle stage (Fig. 3, 4, Table 1). Finally, based on stem basal area, we show that stem growth mainly drives the compositional shift, but the relative contribution of mortality increases with the progression of ecological succession (Fig. 3, 4, Table 1). Thus, in this study, using long-term forest inventory datasets, we demonstrate the link between apparent compositional shifts and changes in the demographic processes during forest succession, in the context of temporal beta diversity.

The patterns of temporal beta diversity have been described occasionally and previous studies specifically reported that the succession rate (i.e., temporal beta diversity) decreases with increased stand age (Shugart & Hett, 1973; Myster & Pickett, 1994; Foster & Tilman, 2000). Moreover, Foster and Tilman (2000) argued the possibility that the succession rate decreases with increasing stand age may be influenced by differences in the life history and longevity of the specific species. The results of this study showed a pattern where the Bray-Curtis dissimilarity index based on stem basal area decreased over time, a finding consistent with previously reported trends (Foster & Tilman, 2000; Anderson, 2007; Li et al., 2016) and one that supports our first hypothesis. As anticipated given our second hypothesis, the relative speed of compositional shifts relative to individual turnover was consistent during the ecological succession for indices based on both stem number (*v*_*s_QL*_) and basal area (*v*_*s_QT*_) turnover (Fig. 2, Table 1). More specifically, the models excluding stand age were selected as the best models for both cases (Table S5). The *v*_*s*_ family of indices were designed to capture changes that take stochastic changes into account, such as individual and basal area turnover (Nakadai, 2020, 2022). This indicates that the contribution of demographic processes (i.e., individual turnover based on both stem number and basal area) to temporal beta diversity (i.e., succession rate) was almost constant across the secondary succession. That is, the succession rate is primarily regulated by individual turnover based on both stem number and basal area. Interestingly, Nakadai (2020) found that *v*_*s_QL*_ values of temperate forests vary with the rate of climate change (i.e., the change rate of annual mean temperature). This result suggests that the contribution of individual turnover based on stem number to apparent compositional shifts (i.e., Bray–Curtis dissimilarity based on stem number) can change with the rate of climate change, ultimately implying that climate change could affect succession rate.

The substitution of species with different life history traits (vital attributes; Noble & Slatyer, 1981) can drive ecological succession in dominance-controlled communities (Foster & Tilman, 2000); in turn, this feeds back to the total changes in the species composition through the demographic processes of each individual (i.e., mortality, growth, and recruitment). Our results clearly demonstrated the relative contributions of the demographic processes to the substitution of species change with succession (Fig. 3, 4, Table 1). At an early stage (i.e., stand initiation phase), species composition changed drastically through recruitment and stem growth. At the intermediate stage (i.e., stem exclusion phase), the composition change was driven by mortality but not by recruitment based on stem number. In the late stages (i.e., understory re-initiation or old-growth phase), the relative contribution of recruitment to the composition change increased again based on stem number, but that of mortality increased based on basal area. These patterns are reasonably consistent with the typical developmental processes of secondary forests (Oliver & Larson, 1996). During the processes, species would be replaced in the order from early successional species to late ones. In fact, Suzuki and Hirao (2018) showed that the community weighted mean in leaf mass per area decreases, suggesting a community-level shade tolerance increase, along with stand age in the same study plots as investigated in this study.

In the analysis based on stem numbers, effects of mean calendar year on the Bray-Curtis dissimilarity index (*d*_*bc*_) and its loss component (*d*_*b*_), the mortality component of individual-based index (*d*_*m*_) and its contribution to apparent compositional shift (*v*_*s_m*_), as well as the total contribution of stem turnover (*v*_*s_mr*_) significantly increased toward recent calendar years (Table 1, S4, Fig. S2); these results imply that species turnover due to mortality tended to increase toward recent years. Although our analyses could not detect the effects of mean calendar year on the mortality component of the individual-based indeces linked to stem basal area (*d*_*M*,_ *v*_*s_M*_), those on the individual-based index (*d*_*MRG*_) and the loss component (*d*_*B*_) were selected based on AIC; indeed, this suggests that species turnover based on basal area, especially species loss, has been increasing in these decades. Suzuki (2021) reported the increase in biomass loss due to stem mortality in the same study plots evaluated here and discussed the potential impact of abiotic and biotic environmental changes such as dry stress, air pollution, and deer herbivory, in these decades. The recent environmental changes can accelerate the species turnover of the secondary forest succession.

The same individual cannot exist in two places at the same time but can exist at one place over time (i.e., at two time steps). This was recently pointed out as a unique feature of temporal beta diversity, more generally, temporal biodiversity changes, compared to spatial studies (Magurran et al., 2019; Nakadai, 2020). This feature has a starting point of individual-based temporal beta diversity which focuses on both individual turnover and persistence (Nakadai, 2020). Approximately half a century ago, in the context of ecological succession, Horn (1975, 1981) focused on the turnover of individuals among different species or within the same species based on the relationship between adult trees and saplings under each adult tree; in addition, the future temporal changes of potential forest compositions were estimated using the Markov chain model. Individual-based temporal beta diversity can partly be recognised as an extended approach of the classical studies focusing on a tree-by-tree replacement process (Horn, 1975, 1981; Korotkov et al., 2001).

The research field of ecological succession originally focused on plants and animals and has recently expanded to various taxonomic groups, including the microbiome (Chuang et al., 2019; Pascual-García & Bell, 2020). Even if the state appears stable in a broad area, ecological successions are always occurring at a certain rate on a smaller spatial scale (Bormann & Likens, 1979). Therefore, because the influences of ecological successions exist ubiquitously, revealing patterns of ecological succession is not only important for understanding those progressive processes but also for understanding the impact of ecological succession on the temporal biodiversity changes at both a regional and global scale. As the sustainable use of natural resources becomes increasingly important, this study clarified the biodiversity changes after anthropogenic disturbance on the aspect of temporal beta diversity. Although we focused on tree communities which are the most classic and central focus of ecological succession research, future studies will need to elucidate patterns at the ecosystem level, including associated organisms.

## Acknowledgments

This study used data from Igarashi et al. (2005), Saito et al. (2014), and Saiki et al. (2019). We appreciate the efforts of all members who contributed to the publication of the data. Ryosuke Nakadai was supported by the Japan Society for the Promotion of Science (22K15188) and the National Institute for Environmental Studies, Japan. Satoshi N. Suzuki was supported by the University of Tokyo.

## Author contributions

Ryosuke Nakadai and Satoshi N. Suzuki conceived and designed the study. Ryosuke Nakadai wrote the first draft and analysed the empirical data with input from Satoshi N. Suzuki. Both authors contributed critically to the drafts, jointly finalised the manuscript, and gave final approval for publication.

## Data Availability statement

Data (in Japanese) was published as the following data papers: Igarashi et al. (2005) (http://doi.org/10.15083/00026215), Saito et al. (2014) (http://doi.org/10.15083/00026149), and Saiki et al. (2019) (https://doi.org/10.15083/00076481). The dataset is available from the Database for the University of Tokyo Forests Experimental and Ecological forest Plots (UTFEEP, http://archives.uf.a.u-tokyo.ac.jp/utfeep/en). The derived datasets are also available from the corresponding author upon reasonable request.

## Supporting information

**Table S1** Census years, plot size, elevation, and year of clear-cutting of tree plots in secondary forests.

**Table S2** List of survey intervals and calculated beta-diversity indices.

**Table S3** Summary of temporal beta-diversity indices used in this study.

**Table S4** Results of a generalised additive mixed model based on the best models selected using an Akaike information criterion (AIC) comparison for temporal beta-diversity indices based on (a) stem number and (b) stem basal area.

**Table S5** Results of the generalised additive mixed model comparison for each index. The selected best models are shown in Table 1 and S4.

**Figure S1** Image showing census year for 15 plots. Star marks indicate the presence of census in the particular years. The numbers on the lines are for counting the number of survey interval.

**Figure S2** Relationship between each component (i.e., gain and loss) of temporal beta-diversity indices and stand age of forest plots situated in a cool temperate secondary forest in Japan. Subplots (a) and (b) depict the indices based on stem number (*d*_*bc*,_ *d*_*b*_ and *d*_*c*_) and basal area (*d*_*BC*_, *d*_*B*_ and *d*_*C*_). Each line was drawn in a situation where all values except stand age were fixed to the mean values for each index. Each coloured area represents the 95% confidence intervals for each index. The model structure and selection details are shown in Table 1 and S5.

## References

Anderson K. J. (2007). Temporal patterns in rates of community change during succession. American Naturalist, 169, 780–793. doi:10.1086/516653

Begon, M., & Townsend, C. R. (2020). Ecology: from individuals to ecosystems. John Wiley & Sons.

Bormann, F. H., & Likens, G. E. (1979). Catastrophic Disturbance and the Steady State in Northern Hardwood Forests: A new look at the role of disturbance in the development of forest ecosystems suggests important implications for land-use policies, American Scientist, 67(6), 660–669.

Brice, M. H., Cazelles, K., Legendre, P., & Fortin, M. J. (2019). Disturbances amplify tree community responses to climate change in the temperate-boreal ecotone. Global Ecology and Biogeography, 28(11), 1668–1681.

Clements, F. E. (1916). Plant succession: an analysis of the development of vegetation (No. 242). Carnegie Institution of Washington.

Chang, C. C., & Turner, B. L. (2019). Ecological succession in a changing world. Journal of Ecology, 107(2), 503–509.

Chazdon R. L., Letcher S. G., Van Breugel M., Martínez-Ramos M., Bongers F., & Finegan B. (2007). Rates of change in tree communities of secondary Neotropical forests following major disturbances. Philosophical Transactions of the Royal Society B: Biological Sciences, 362, 273–289. doi:10.1098/rstb.2006.1990

Chuang, J. S., Frentz, Z., & Leibler, S. (2019). Homeorhesis and ecological succession quantified in synthetic microbial ecosystems. Proceedings of the National Academy of Sciences, 116(30), 14852–14861.

Foster, B. L., & Tilman, D. (2000). Dynamic and static views of succession: testing the descriptive power of the chronosequence approach. Plant Ecology, 146(1), 1–10.

Fregin, S., Haase, M., Olsson, U., & Alström, P. (2012). Pitfalls in comparisons of genetic distances: a case study of the avian family Acrocephalidae. Molecular Phylogenetics and Evolution, 62(1), 319–328.

Gotelli, N. J., Moyes, F., Antäo, L. H., Blowes, S. A., Dornelas, M., McGill, B. J., & Magurran, A. E. (2022). Long[term changes in temperate marine fish assemblages are driven by a small subset of species. Global Change Biology, 28(1), 46–53.

Hill, M. F., Witman, J. D., & Caswell, H. (2002). Spatio-temporal variation in Markov chain models of subtidal community succession. Ecology Letters, 5(5), 665–675.

Horn, H. S. (1975). Markovian properties of forest succession. Ecology and Evolution of Communities, 196–211.

Horn, H. S. (1981). Succession. Theoretical Ecology. Ecology and Evolution of Communities, 187–204.

Igarashi, Y., Oomura, K., & Fujiwara, A. (2005). Growth records on the secondary forest permanent plots in University Forest in Chichibu. Miscellaneous Information, The University of Tokyo Forests, 44, 121–210. (in Japanese).

Inouye, R. S., Allison, T. D., & Johnson, N. C. (1994). Old field succession on a Minnesota sand plain: effects of deer and other factors on invasion by trees. Bulletin of the Torrey Botanical Club, 266–276.

Korotkov, V. N., Logofet, D. O., & Loreau, M. (2001). Succession in mixed boreal forest of Russia: Markov models and non-Markov effects. Ecological Modelling, 142(1–2), 25–38.

Legendre, P. (2019). A temporal beta-diversity index to identify sites that have changed in exceptional ways in space–time surveys. Ecology and Evolution, 9, 3500–3514.

Li, S. P., Cadotte, M. W., Meiners, S. J., Pu, Z., Fukami, T., & Jiang, L. (2016). Convergence and divergence in a long-term old-field succession: The importance of spatial scale and species abundance. Ecology Letters, 19(9), 1101–1109.

Magurran, A. E., Dornelas, M., Moyes, F., & Henderson, P. A. (2019). Temporal β diversity: A macroecological perspective. Global Ecology and Biogeography, 28, 1949–1960.

Myster R. W., & Pickett S. T. A. (1994). A comparison of rate of succession over 18 yr in 10 contrasting old fields. Ecology, 75, 387–392.

Nakadai, R. (2020). Degrees of compositional shift in tree communities vary along a gradient of temperature change rates over one decade: Application of an individual-based temporal beta-diversity concept. Ecology and Evolution, 10(24), 13613–13623.

Nakadai, R. (2021). Individual-based multiple-unit dissimilarity: novel indices and null model for assessing temporal variability in community composition. Oecologia, 197(2), 353–364.

Nakadai, R. (2022). Development of novel temporal beta-diversity indices for assessing community compositional shifts accounting for changes in the properties of individuals. Ecological indicators. doi: 10.1016/j.ecolind.2022.109427

Noble, I. R., & Slatyer, R. O. (1981). Concepts and models of succession in vascular plant communities subject to recurrent fire. In Conference on Fire and the Australian Biota, Canberra (Australia), 9 Oct 1978. Australian Academy of Science.

Odum, E. P. (1950). Bird populations of the highlands (North Carolina) plateau in relation to plant succession and avian invasion. Ecology 31:587–605

Oliver C. H., & Larson B. C. (1996). Forest Stand Dynamics, update edition, John Wiley & Sons. Inc.

Prach, K., & Walker, L. R. (2011). Four opportunities for studies of ecological succession. Trends in Ecology & Evolution, 26, 119–123.

Pascual-García, A., & Bell, T. (2020). Community-level signatures of ecological succession in natural bacterial communities. Nature communications, 11(1), 1–11.

Pickett, S. T. (1989). Space-for-time substitution as an alternative to long-term studies. In Long-term studies in ecology (pp. 110–135). Springer, New York, NY.

Rees, M., Condit, R., Crawley, M., Pacala, S., & Tilman, D. (2001). Long-term studies of vegetation dynamics. Science, 293(5530), 650–655.

Saiki, M., Takatoku, K., Igarashi, Y., & Haraguchi, R. (2019). Growth Records on the Secondary Forest Permanent Plots in the University of Tokyo Chichibu Forest (2017). Miscellaneous Information, The University of Tokyo Forests, 61, 19–25. (in Japanese).

Saito, T., Saiki, M., Aikawa, M., & Kurita, N. (2014). Growth records on the secondary forest permanent plots in the University of Tokyo Forest (2007, 2012).

Miscellaneous Information, The University of Tokyo Forests, 56, 197–286. (in Japanese).

Shimadzu, H. et al. (2015). Measuring temporal turnover in ecological communities. Methods in Ecology and Evolution, 6, 1384–1394.

Shugart, H. H., & Hett, J. M. (1973). Succession: similarities of species turnover rates. Science, 180(4093), 1379–1381.

Suzuki, S. N., & Hirao, T. (2018). Recruitment drives successional changes in the community[level leaf mass per area in a winter[deciduous broad[leaf forest. Journal of Vegetation Science, 29(4), 756–764.

Suzuki, S. N. (2021). Acceleration and deceleration of aboveground biomass accumulation rate in a temperate forest in central Japan. Forest Ecology and Management, 479, 118550.

Tatsumi, S., Iritani, R., & Cadotte, M. W. (2021). Temporal changes in spatial variation: Partitioning the extinction and colonisation components of beta diversity. Ecology Letters, 24(5), 1063–1072.

Yodzis, P. (1986). Competition, mortality, and community structure, in “Community Ecology” (J. Diamond and TJ Case, Eds.).

