## Supplementary material for "Ecological succession revisited from a temporal beta-diversity perspective": Table S3

Table S3. Summary of temporal beta-diversity indices used in this study.

| Index | Formula | Reference | Ecological interpretation |
| --- | --- | --- | --- |
| (a) Temporal beta-diversity indices based on stem number | |  |  |
| *d_bc_* | $\frac{b+c}{2a+b+c}= \frac{\sum_{i} \left[ x_{ij}-min(x_{ij},x_{ik}) \right]+\sum_{i} \left[ x_{ik}-min(x_{ij},x_{ik}) \right]}{2[\sum_{i} min(x_{ij},x_{ik})]+\sum_{i} \left[ x_{ij}-min(x_{ij},x_{ik}) \right]+\sum_{i} \left[ x_{ik}-min(x_{ij},x_{ik}) \right]}$ | Odum (1950) | Apparent compositional shift based on stem number. |
| *d_mr_* | $\frac{m+r}{2p+m+r}=\frac{\sum_{o} \left[ z_{oj}-min(z_{oj},z_{ok}) \right]+\sum_{o} \left[ z_{ok}-min(z_{oj},z_{ok}) \right]}{2[\sum_{o} min(z_{oj},z_{ok})]+\sum_{o} \left[ z_{oj}-min(z_{oj},z_{ok}) \right]+\sum_{o} \left[ z_{ok}-min(z_{oj},z_{ok}) \right]}$ | Nakadai (2020) | Total turnover based on stem number. |
| *v_s.QL_* | $\frac{b+c}{2e+b+c}=\frac{b+c}{m+r}= \frac{\sum_{i} \left[ x_{ij}-min(x_{ij},x_{ik}) \right]+\sum_{i} \left[ x_{ik}-min(x_{ij},x_{ik}) \right]}{2[\sum_{i} min(x_{ij},x_{ik})- \sum_{o} min(z_{oj},z_{ok})]+\sum_{i} \left[ x_{ij}-min(x_{ij},x_{ik}) \right]+\sum_{i} \left[ x_{ik}-min(x_{ij},x_{ik}) \right]}$ | Nakadai (2020) | Ratio of compositional shift against turnover based on stem number. |
| *d_b_* | $\frac{b}{2a+b+c}= \frac{\sum_{i} \left[ x_{ij}-min(x_{ij},x_{ik}) \right]}{2[\sum_{i} min(x_{ij},x_{ik})]+\sum_{i} \left[ x_{ij}-min(x_{ij},x_{ik}) \right]+\sum_{i} \left[ x_{ik}-min(x_{ij},x_{ik}) \right]}$ | Legendre (2019) | Contribution of apparent compositional loss in *d_bc_.* |
| *d_c_* | $\frac{c}{2a+b+c}= \frac{\sum_{i} \left[ x_{ik}-min(x_{ij},x_{ik}) \right]}{2[\sum_{i} min(x_{ij},x_{ik})]+\sum_{i} \left[ x_{ij}-min(x_{ij},x_{ik}) \right]+\sum_{i} \left[ x_{ik}-min(x_{ij},x_{ik}) \right]}$ | Legendre (2019) | Contribution of apparent compositional gain in *d_bc_.* |
| *d_m_* | $\frac{m}{2p+m+r}=\frac{\sum_{o} \left[ z_{oj}-min(z_{oj},z_{ok}) \right]}{2[\sum_{o} min(z_{oj},z_{ok})]+\sum_{o} \left[ z_{oj}-min(z_{oj},z_{ok}) \right]+\sum_{o} \left[ z_{ok}-min(z_{oj},z_{ok}) \right]}$ | Nakadai (2020) | Contribution of mortal individuals in *d_mr_.* |
| *d_r_* | $\frac{r}{2p+m+r}=\frac{\sum_{o} \left[ z_{ok}-min(z_{oj},z_{ok}) \right]}{2[\sum_{o} min(z_{oj},z_{ok})]+\sum_{o} \left[ z_{oj}-min(z_{oj},z_{ok}) \right]+\sum_{o} \left[ z_{ok}-min(z_{oj},z_{ok}) \right]}$ | Nakadai (2020) | Contribution of recruit individuals in *d_mr_.* |
| *v_s_mr_* | $v_{s.mr}=d_{mr}\times v_{s.QL}$ | Nakadai (2020)* | Total contribution of turnover based on stem number to apparent compositional shift. |
| *v_s_m_* | $v_{s.m}=d_{m}\times v_{s.QL}$ | This study | Contribution of mortality in *v_s_mr_.* |
| *v_s_r_* | $v_{s.r}=d_{r}\times v_{s.QL}$ | This study | Contribution of recruitment in *v_s_mr_.* |
| (b) Temporal beta-diversity indices based on stem basal area | |  |  |
| *d_BC_* | $\frac{B+C}{2A+B+C}= \frac{\sum_{i} \left[ y_{ij}-min(y_{ij},y_{ik}) \right]+\sum_{i} \left[ y_{ik}-min(y_{ij},y_{ik}) \right]}{2[\sum_{i} min(y_{ij},y_{ik})]+\sum_{i} \left[ y_{ij}-min(y_{ij},y_{ik}) \right]+\sum_{i} \left[ y_{ik}-min(y_{ij},y_{ik}) \right]}$ | Odum (1950) | Apparent compositional shift based on stem basal area. |
| *d_MRG_* | $\frac{M+R+G}{2P+M+R+G}= \frac{\sum_{o} \left[ \left[ w_{oj}-min(w_{oj},w_{ok}) \right]\times\left[ z_{oj}-min(z_{oj},z_{ok}) \right] \right]+\sum_{o} \left[ \left[ w_{ok}-\min\left( w_{oj},w_{ok} \right) \right]\times\left[ z_{ok}-min(z_{oj},z_{ok}) \right] \right]+\sum_{o} \left[ \left[ \max\left( w_{oj},w_{ok} \right)-\min\left( w_{oj},w_{ok} \right) \right]\times min(z_{oj},z_{ok}) \right]}{2[\sum_{o} min(w_{oj},w_{ok})+\sum_{o} \left[ \left[ w_{oj}-min(w_{oj},w_{ok}) \right]\times\left[ z_{oj}-min(z_{oj},z_{ok}) \right] \right]+\sum_{o} \left[ \left[ w_{ok}-\min\left( w_{oj},w_{ok} \right) \right]\times\left[ z_{ok}-min(z_{oj},z_{ok}) \right] \right]+\sum_{o} \left[ \left[ \max\left( w_{oj},w_{ok} \right)-\min\left( w_{oj},w_{ok} \right) \right]\times min(z_{oj},z_{ok}) \right]}$ | Nakadai (2022) | Total turnover based on stem number. |
| *v_s.QT_* | $\frac{B+C}{2E+B+C}=\frac{B+C}{M+R}= \frac{\sum_{i} \left[ y_{ij}-min(y_{ij},y_{ik}) \right]+\sum_{i} \left[ y_{ik}-min(y_{ij},y_{ik}) \right]}{2[\sum_{i} min(y_{ij},y_{ik})- \sum_{o} min(w_{oj},w_{ok})]+\sum_{i} \left[ y_{ij}-min(y_{ij},y_{ik}) \right]+\sum_{i} \left[ y_{ik}-min(y_{ij},y_{ik}) \right]}$ | Nakadai (2022) | Ratio of compositional shift against turnover based on stem basal area. |
| *d_B_* | $\frac{B}{2A+B+C}= \frac{\sum_{i} \left[ y_{ij}-min(y_{ij},y_{ik}) \right]}{2[\sum_{i} min(y_{ij},y_{ik})]+\sum_{i} \left[ y_{ij}-min(y_{ij},y_{ik}) \right]+\sum_{i} \left[ y_{ik}-min(y_{ij},y_{ik}) \right]}$ | Legendre (2019) | Contribution of apparent compositional loss in *d_BC_.* |
| *d_C_* | $\frac{C}{2A+B+C}= \frac{\sum_{i} \left[ y_{ik}-min(y_{ij},y_{ik}) \right]}{2[\sum_{i} min(y_{ij},y_{ik})]+\sum_{i} \left[ y_{ij}-min(y_{ij},y_{ik}) \right]+\sum_{i} \left[ y_{ik}-min(y_{ij},y_{ik}) \right]}$ | Legendre (2019) | Contribution of apparent compositional gain in *d_BC_.* |
| *d_M_* | $\frac{M}{2P+M+R+G}= \frac{\sum_{o} \left[ \left[ w_{oj}-min(w_{oj},w_{ok}) \right]\times\left[ z_{oj}-min(z_{oj},z_{ok}) \right] \right]}{2[\sum_{o} min(w_{oj},w_{ok})+\sum_{o} \left[ \left[ w_{oj}-min(w_{oj},w_{ok}) \right]\times\left[ z_{oj}-min(z_{oj},z_{ok}) \right] \right]+\sum_{o} \left[ \left[ w_{ok}-\min\left( w_{oj},w_{ok} \right) \right]\times\left[ z_{ok}-min(z_{oj},z_{ok}) \right] \right]+\sum_{o} \left[ \left[ \max\left( w_{oj},w_{ok} \right)-\min\left( w_{oj},w_{ok} \right) \right]\times min(z_{oj},z_{ok}) \right]}$ | Nakadai (2022) | Contribution of mortal individuals in *d_MRG_.* |
| *d_R_* | $\frac{R}{2P+M+R+G}= \frac{\sum_{o} \left[ \left[ w_{ok}-\min\left( w_{oj},w_{ok} \right) \right]\times\left[ z_{ok}-min(z_{oj},z_{ok}) \right] \right]}{2[\sum_{o} min(w_{oj},w_{ok})+\sum_{o} \left[ \left[ w_{oj}-min(w_{oj},w_{ok}) \right]\times\left[ z_{oj}-min(z_{oj},z_{ok}) \right] \right]+\sum_{o} \left[ \left[ w_{ok}-\min\left( w_{oj},w_{ok} \right) \right]\times\left[ z_{ok}-min(z_{oj},z_{ok}) \right] \right]+\sum_{o} \left[ \left[ \max\left( w_{oj},w_{ok} \right)-\min\left( w_{oj},w_{ok} \right) \right]\times min(z_{oj},z_{ok}) \right]}$ | Nakadai (2022) | Contribution of recruited individuals in *d_MRG_.* |
| *d_G_* | $\frac{G}{2P+M+R+G}= \frac{\sum_{o} \left[ \left[ \max\left( w_{oj},w_{ok} \right)-\min\left( w_{oj},w_{ok} \right) \right]\times min(z_{oj},z_{ok}) \right]}{2[\sum_{o} min(w_{oj},w_{ok})+\sum_{o} \left[ \left[ w_{oj}-min(w_{oj},w_{ok}) \right]\times\left[ z_{oj}-min(z_{oj},z_{ok}) \right] \right]+\sum_{o} \left[ \left[ w_{ok}-\min\left( w_{oj},w_{ok} \right) \right]\times\left[ z_{ok}-min(z_{oj},z_{ok}) \right] \right]+\sum_{o} \left[ \left[ \max\left( w_{oj},w_{ok} \right)-\min\left( w_{oj},w_{ok} \right) \right]\times min(z_{oj},z_{ok}) \right]}$ | Nakadai (2022) | Contribution of individual growth in *d_MRG_.* |
| *v_s_MRG_* | $v_{s.MRG}=d_{MRG}\times v_{s.QT}$ | This study | Total contribution of turnover based on stem basal area to apparent compositional shift. |
| *v_s_M_* | $v_{s.M}=d_{M}\times v_{s.QT}$ | This study | Contribution of mortal individuals in *v_s_MRG_.* |
| *v_s_R_* | $v_{s.R}=d_{R}\times v_{s.QT}$ | This study | Contribution of recruited individuals in *v_s_MRG_.* |
| *v_s_G_* | $v_{s.G}=d_{G}\times v_{s.QT}$ | This study | Contribution of individual growth in *v_s_MRG_.* |

*x_ij_* is the stem number of species *i* at temporal unit *j*, and *x_ik_* is the stem number of species *i* at temporal unit *k*. *y_ij_* is the stem basal area of species *i* at time T*j* at a site, and *y_ik_* is the stem basal area of species *i* at time T*k* at a site. *z_oj_* is individual *o* at time T*j*, and *z_ok_* is individual *o* at time T*k*. Both *z_oj_* and *z_ok_* have values of one or zero (i.e., presence and absence, respectively). *w_oj_* is the measure of individual *o* at time T*j*, and *w_ok_* is the measure of individual *o* at time T*k*. *m* refers to the number of individuals in existence at T*j* but are dead prior to T*k*, thereby separately identifying the deaths of individuals between T*j* and T*k*; *r* refers to the number of individuals not yet living at T*j* but are present at T*k*, thereby identifying the recruitment of individuals between T*j* and T*k*; *p* refers to the number of individuals that persisted from T*j* to T*k*. *e* refers to the number of individuals that did not contribute to the apparent compositional shift among mortality or recruitment individuals (please see Nakadai 2020 for further details). In the components based on stem basal area, *M* refers to the total basal area of individuals occurring at T*j* that die prior to T*k*, thereby representing the total basal area of the mortal individuals between T*j* and T*k*; *R* refers to the total basal area of individuals not yet living at T*j* but are alive and counted at T*k*, thereby representing the individuals recruited between T*j* and T*k*. For persistent individuals, the original and altered total basal area are partitioned into persistent (*P*) and growth (*G*) components, respectively. *E* refers to the total basal area that did not contribute to the apparent compositional shift among mortality, recruitment, or growth of individuals (please see Nakadai 2022 for further details).

*The index *v_s_mr_* is identical to the contribution of compositional shift (*d_s_*=(b+c)/(2p+m+r)) introduced in Nakadai (2020).
