## Supplementary figures and images for "Ecological succession revisited from a temporal beta-diversity perspective"

### Figure S1

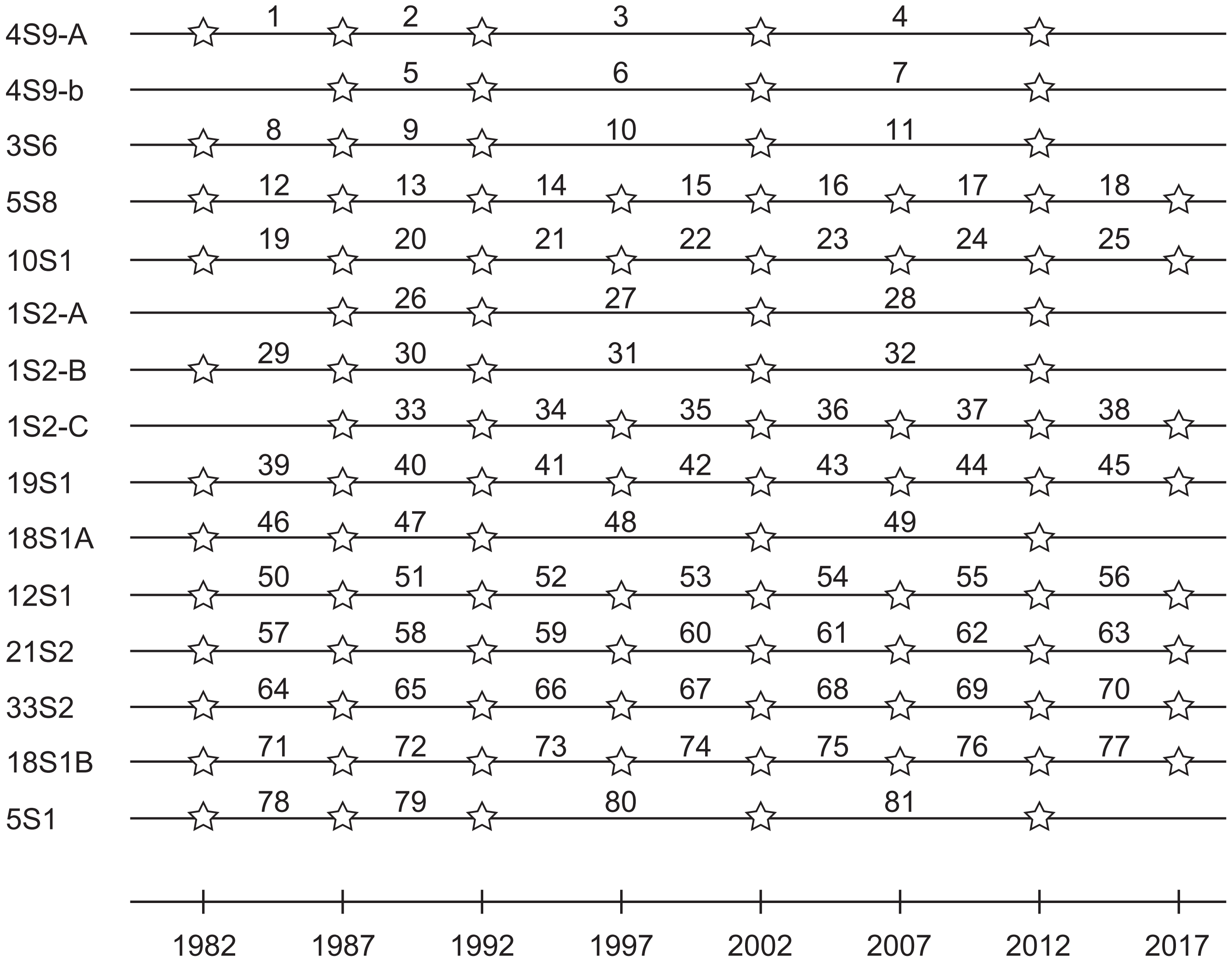

### Figure S2

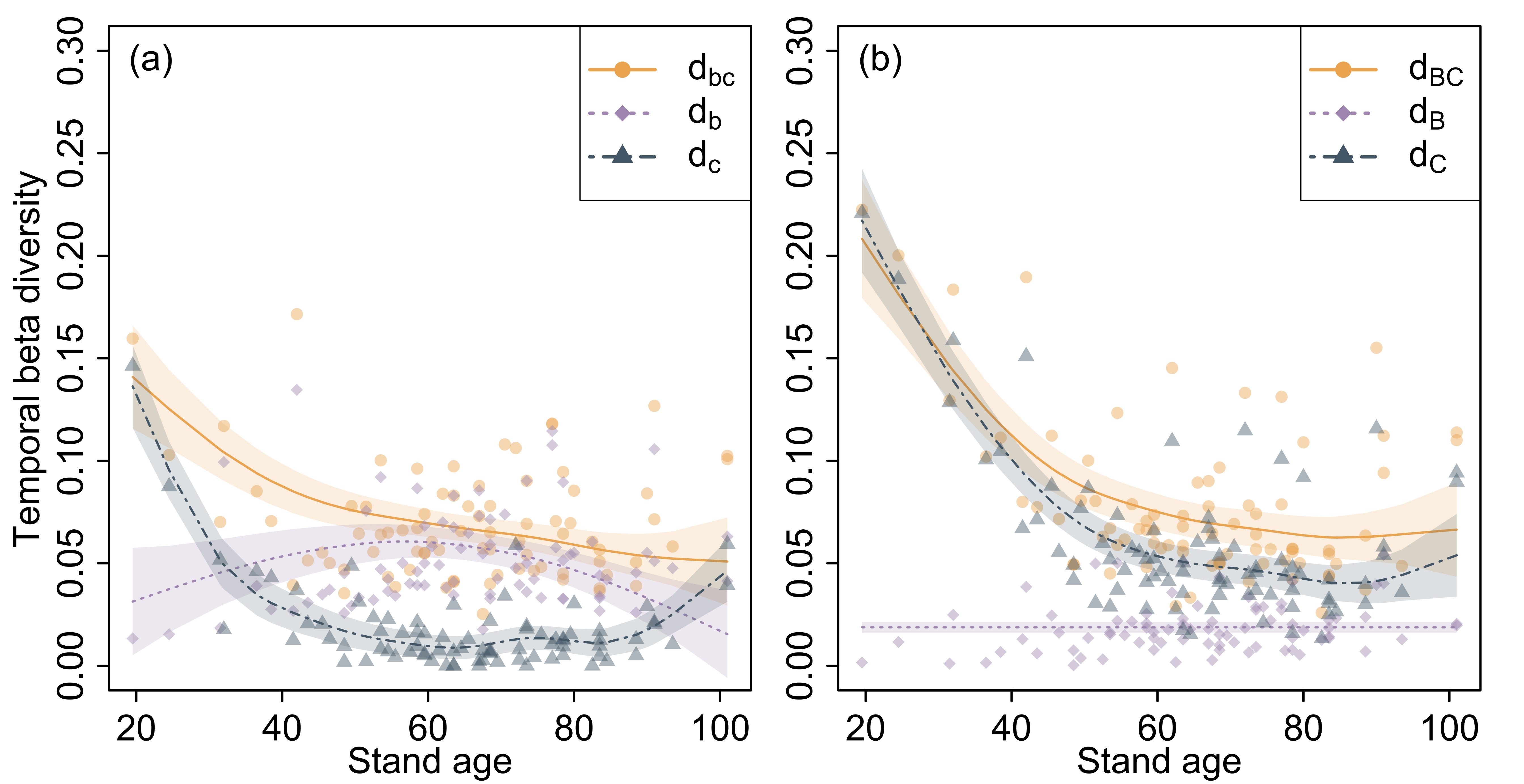
